# Embryonic development in the acoel *Hofstenia miamia*

**DOI:** 10.1101/2021.01.25.427674

**Authors:** Julian O. Kimura, Lorenzo Ricci, Mansi Srivastava

## Abstract

Acoels are marine worms that belong to the phylum Xenacoelomorpha. The phylogenetic placement of this group as a deep-diverging lineage makes acoel embryos an attractive system to study the evolution of major bilaterian traits. Thus far, acoel development has not been described in detail at the morphological and transcriptomic levels in a species where functional genetic studies are possible. Here, we present a set of developmental landmarks for embryogenesis in the highly regenerative acoel *Hofstenia miamia*. We generated a developmental staging atlas from zygote to hatched worm based on gross morphology, with accompanying bulk transcriptome data for each of the stages. *Hofstenia* embryos undergo a stereotyped cleavage program known as duet cleavage, which results in two large ‘macromeres’ at the vegetal pole and numerous small ‘micromeres’ at the animal pole. The macromeres become internalized as micromere progeny proliferate and move vegetally, enveloping the larger blastomeres. We also noted a second, previously undescribed cell internalization event at the animal pole, following which we detected tissues corresponding to all three germ layers. Our work on *Hofstenia* embryos provides a resource for future investigations of acoel development, which will yield insights into the evolution of development and regeneration.

**Summary Statement:** Comprehensive characterization of embryonic development in the acoel worm *Hofstenia miamia* with accompanying transcriptome data.

## Introduction

Acoel worms are a group of marine invertebrates that have garnered attention in the fields of evolutionary and regenerative biology due to their phylogenetic placement and the extensive regenerative capacity that some species possess (Bourlat and Hejnol, 2009; Gehrke et al., 2019; Gehrke and Srivastava, 2016; Hejnol and Pang, 2016; Srivastava et al., 2014). Based on morphological characteristics and striking similarities in cell cleavage patterns during early embryogenesis, acoels were thought to belong to the phylum Platyhelminthes (Ax and Dörjes, 1966; Ax and Jeffries, 1987; Boyer et al., 1996; Boyer and Jonathan, 1998; Bresslau, 1909; Costello and Henley, 1976; Henry and Martindale, 1999; Hyman, n.d.; Peterson and Eernisse, 2001; Smith et al., 1986). Members of the phylum Platyhelminthes develop using an ancestral cleavage program called spiral cleavage (four large vegetal blastomeres producing smaller cells toward the animal pole), whereas acoel worms undergo duet cleavage (two large vegetal blastomeres producing smaller cells toward the animal pole) (Apelt, 1969; Boyer, 1971; Bresslau, 1909; Henry et al., 2000; Maslakova et al., 2004; Nielsen, 1995). Given that acoels were nested within a group with an ancestral spiral cleavage program, it was hypothesized that duet cleavage was a derived form of spiral cleavage. However, recent molecular phylogenetic analyses have revealed that acoels belong to the major animal clade Xenacoelomorpha, which represents either the sister group to all other bilaterians (Nephrozoa) or to a deuterostome lineage (Ambulacraria) (Fig. 1A) (Hejnol et al., 2009; Jondelius et al., 2011; Kapli and Telford, 2020; Marlétaz et al., 2019; Mwinyi et al., 2010; Philippe et al., 2007, 2011, 2019; Ruiz-Trillo et al., 1999, 2002, 2004; Ruiz-Trillo and Paps, 2016; Sempere et al., 2007; Telford et al., 2003). Both of these phylogenetic positions for Xenacoelomorpha make acoels highly informative for understanding the evolution of bilaterian traits. Moreover, in either scenario, xenacoelomorphs are distantly-related to Platyhelminthes and to other animals that undergo spiral cleavage. This raises the possibility that the cleavage program in acoels represents an independently-evolved yet understudied mode of development. Therefore, in addition to providing insights into the evolution of bilateral symmetry, mesoderm, and centralized nervous system (Bourlat and Hejnol, 2009; Hejnol and Pang, 2016), studies of acoel embryonic development could reveal new mechanisms of development.

**Fig. 1.**
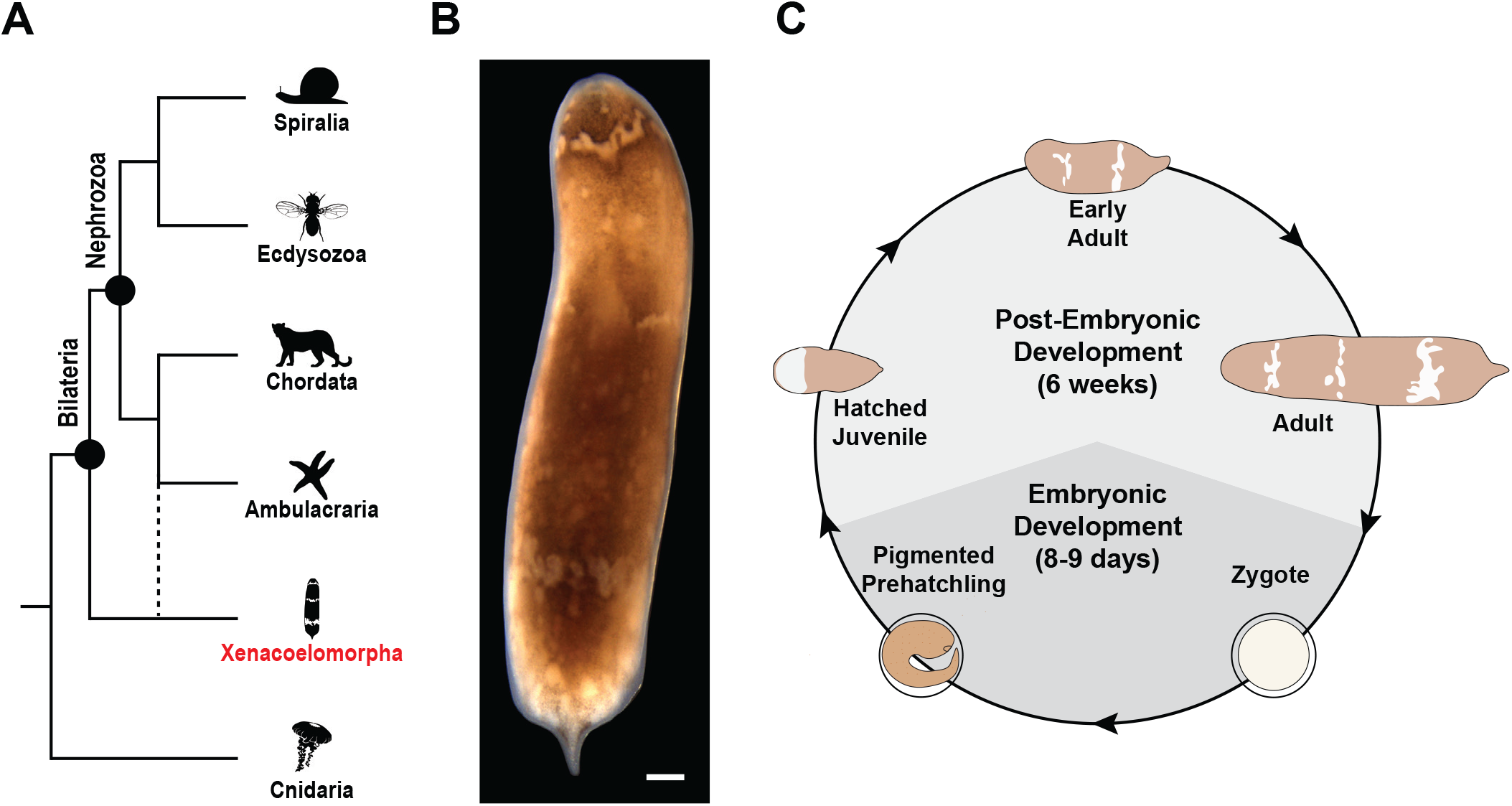
*Hofstenia miamia* as an acoel model for studies of embryogenesis. (A) Acoels belong to the phylum Xenacoelomorpha (red) that is either sister to all other bilaterians (nephrozoans) or to ambulacrarians. (B) Brightfield image of an adult *Hofstenia miamia.* (C) *Hofstenia miamia* life cycle, based on a schematic developed by Dr. Yi Jyun Luo. Light-gray represents post-embryonic development, while dark-gray represents embryonic development. Sexually mature adult worms lay fertilized zygotes that hatch into a Hatched Juvenile worm in 8-9 days. The Hatched Juvenile worm then undergoes post embryonic development into a sexually mature adult. Scale bar, 100μm.

Various aspects of acoel development have been described via studies in different species (Apelt, 1969; Boyer, 1971; Bresslau, 1909; Hejnol and Martindale, 2008a, 2008b; Henry et al., 2000; Ladurner and Rieger, 2000; Perea-Atienza et al., 2018; Ramachandra et al., 2002; Semmler et al., 2008). However, a comprehensive description from zygote to hatching is lacking in an acoel model where functional genetic studies are possible. All acoel species studied thus far were found to undergo duet cleavage (Apelt, 1969; Boyer, 1971; Bresslau, 1909; Henry et al., 2000), which features embryos with two large blastomeres called macromeres on the vegetal pole that divide asymmetrically to produce several smaller blastomeres called micromeres on the animal pole (Fig. S1A). The large, vegetal macromeres are then internalized as the micromeres continue to proliferate and move towards the vegetal pole. Fate-mapping experiments in the species *Neochildia fusca* revealed that micromeres give rise to ectodermal fates, whereas macromeres labeled just prior to internalization give rise to the endomesoderm (Henry et al., 2000). Furthermore, the expression of the conserved blastopore marker *brachyury* was detected at the site of macromere internalization in the species *Convolutriloba longifissura* (Hejnol and Martindale, 2008a). Although these results are from two separate species, given the conservation of the duet cleavage program, it was hypothesized that the process of macromere internalization represented gastrulation in acoels. Insights into acoel development such as staging, gene expression, and myogenesis are available from different species (Hejnol and Martindale, 2008a, 2008b; Ladurner and Rieger, 2000; Perea-Atienza et al., 2018; Ramachandra et al., 2002; Semmler et al., 2008). However, systematic studies of embryogenesis in one system, particularly a genetically tractable research organism, are needed to obtain mechanistic insights into acoel development and the evolution of bilaterian traits.

Here, we present a morphological and molecular characterization of embryogenesis in the acoel worm *Hofstenia miamia*, a new research organism for studying acoel development and regeneration (Fig. 1B,C) (Gehrke et al., 2019; Srivastava et al., 2014). *Hofstenia* is a genetically tractable model system with molecular resources including a high quality genome and transcriptome (Gehrke et al., 2019; Srivastava et al., 2014). *Hofstenia* can be cultured easily in the lab and produces plentiful, accessible embryos that are amenable to experimental manipulation (Fig. S1B,C). Furthermore, *Hofstenia*’s ability to undergo whole-body regeneration using pluripotent stem cells allows for the unique opportunity to study how a highly regenerative animal undergoes development. *Hofstenia*’s established repertoire of tools, such as systemic RNAi, for studying regeneration makes it an excellent model for studying acoel development.

We staged *Hofstenia* embryos from zygote to hatching over a nine day period, characterized their duet cleavage pattern, identified two major cell internalization events, generated transcriptome data in bulk for each stage, and determined the timing for the detection of differentiated tissues. This work serves as a foundation for future mechanistic studies of acoel embryogenesis and regeneration, which could ultimately provide insights into the evolution of bilaterian traits and regeneration.

## Results

### A developmental atlas for Hofstenia miamia

We generated a developmental atlas based on gross morphology, phalloidin staining, and nuclear labeling for *Hofstenia miamia* (Fig. 2A,B; Fig. 3A,B; and Movie S1). Developmental timing is denoted in hours post laying (hpl) at 23°C. Hermaphroditic adults produce embryos that develop directly into juvenile worms that hatched in eight to nine days (Fig. 1C). The variability in hatching time appears to be attributed to the time it takes for the animal to break through its egg shell rather than the timing of developmental milestones. Embryos are laid in clutches as spherical, fertilized zygotes (0hpl), about 300μm in diameter (Fig. 2A; Fig. S1D; Movie S1; and Movie S2). The zygotes are opaque, each enveloped in a clear egg shell upon laying.

**Fig. 2.**
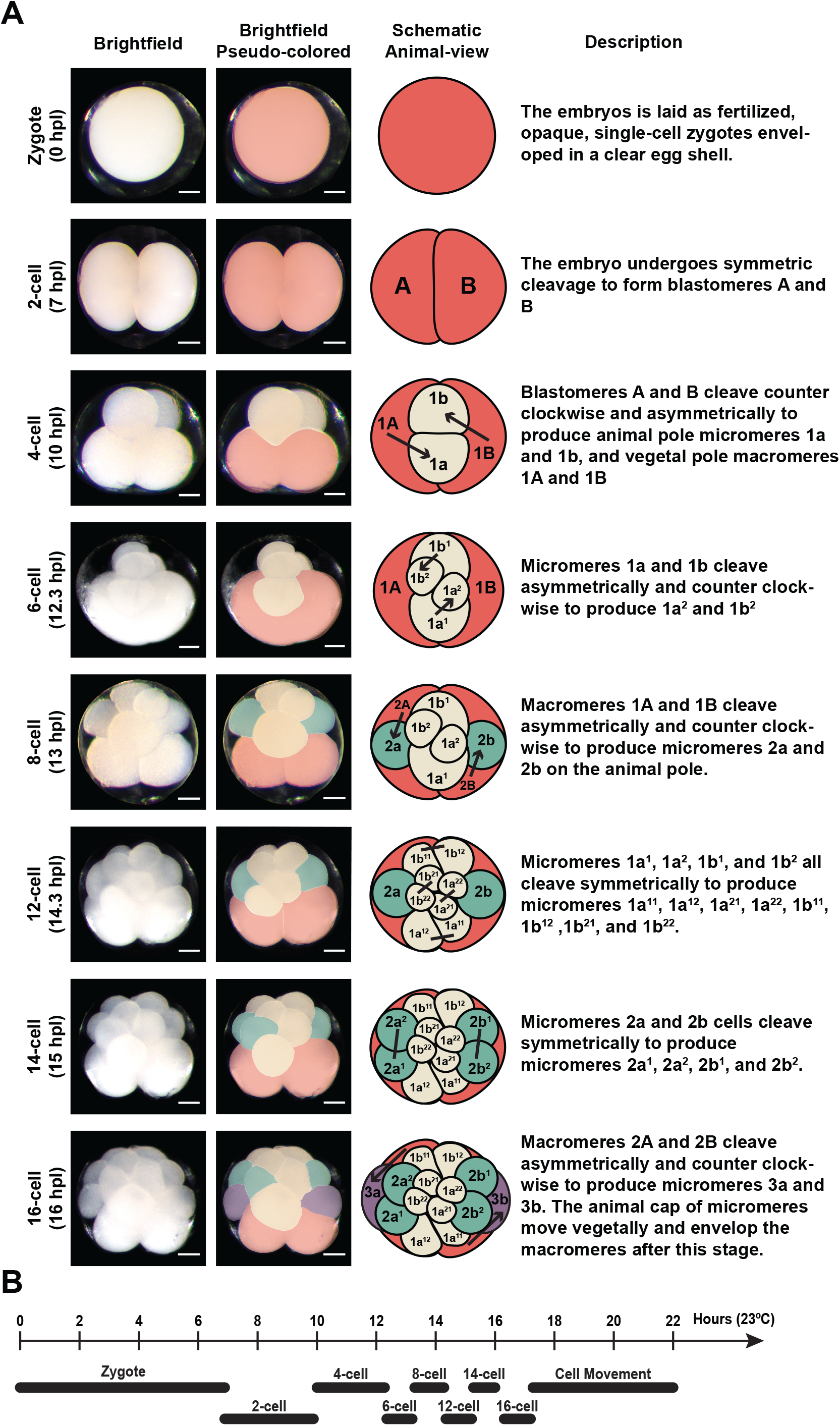
The acoel worm *Hofstenia miamia* undergoes a stereotyped, duet cleavage program. (A) From left to right: developmental atlas of early cleavage showing representative brightfield images of live *Hofstenia* embryos; representative brightfield images pseudo-colored to show distinct blastomeres and their daughter cells; animal-view schematic representation of the cleavage order with arrows signifying asymmetric cleavage and lines signifying symmetric cleavage; defining characteristics of each developmental stage. Red: macromeres; yellow: first set of micromeres and their progeny; green: second set of micromeres and their progeny; purple: third set of micromeres. (B) Schematic timeline of *Hofstenia* early cleavage. Scale bars, 100μm.

**Fig. 3.**
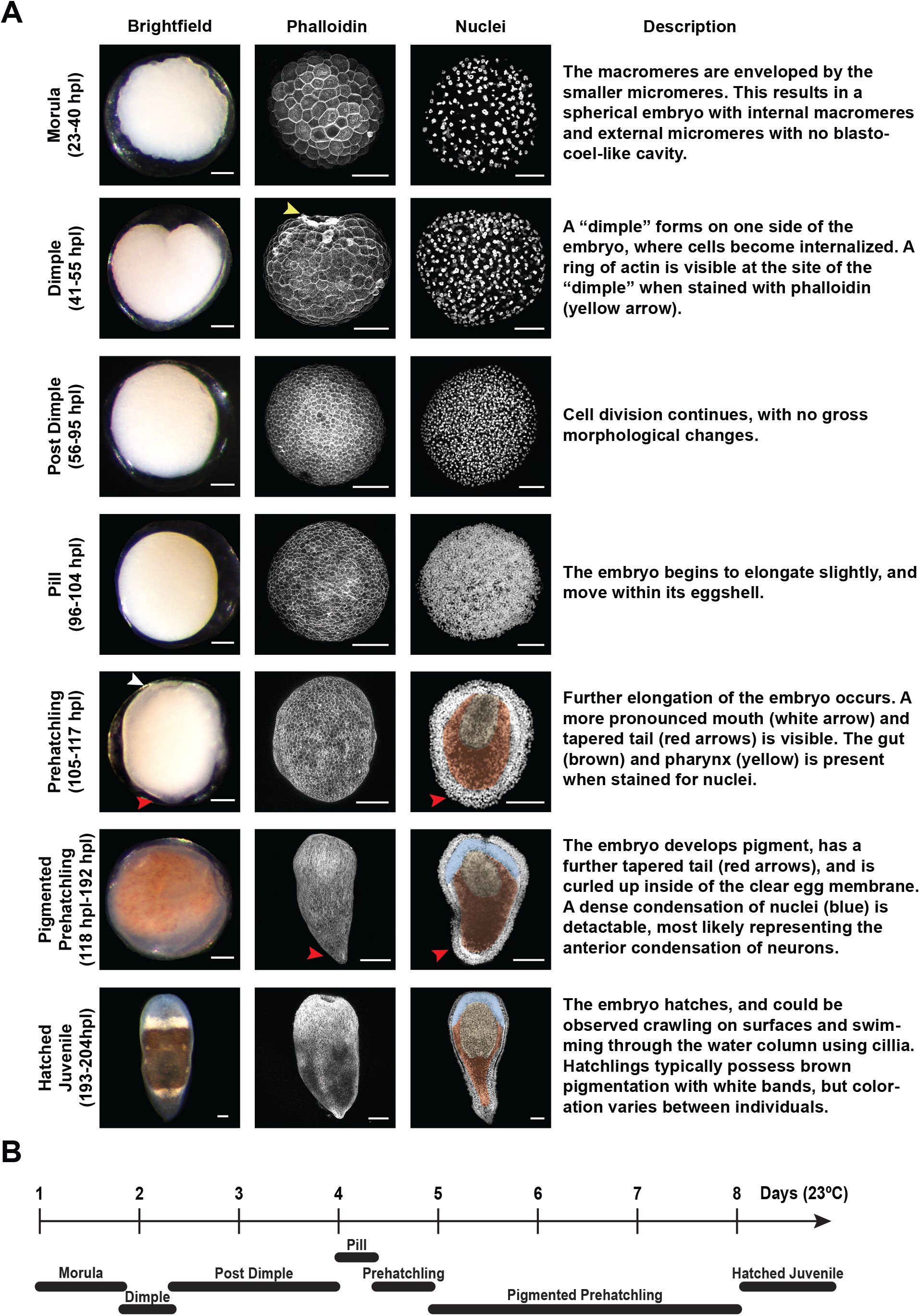
Developmental atlas of later stages during *Hofstenia miamia* embryogenesis. (A) From left to right: developmental atlas of later stages of *Hofstenia* embryogenesis showing representative brightfield images; phalloidin and nuclear staining of each corresponding developmental stage; defining characteristics of each developmental stage. Yellow Arrow: Ring of actin at the site of the “dimple”; White arrow: Opening of the mouth; Red Arrows: Tapered posterior; Yellow Shading: Pharynx; Brown Shading: Gut; Blue Shading: Anterior condensation of neurons. (B) Schematic timeline of later *Hofstenia* developmental stages. Scale bars, 100μm.

### Early Cleavage Stage (0 hpl - 22 hpl)

Much like previously-studied acoels, *Hofstenia* embryos undergo a stereotyped, duet cleavage pattern during early development. The Early Cleavage stage starts with the initiation of cell division, and ends with a cell movement event that results in the internalization of vegetal cells (Fig. 2A; Fig. S2A; Movie S1; and Movie S2). The first cleavage is symmetric, and produces two blastomeres that we refer to as A and B (7 hpl), following the convention in previously-published descriptions of acoel embryogenesis (Boyer, 1971; Bresslau, 1909; Henry et al., 2000). During second cleavage, blastomeres A and B divide synchronously and asymmetrically, producing smaller progeny towards the animal pole laeotropically, *i.e.*, in a counter-clockwise orientation when viewed from the animal pole. This results in a 4-cell embryo where smaller cells, micromeres 1a and 1b, are generated at a 45° angle to the animal-vegetal (AV) axis. The larger cells on the vegetal pole are now referred to as macromeres 1A and 1B. Micromeres 1a and 1b sit at the cell junction of macromeres 1A and 1B.

Next, micromeres and macromeres enter a phase of sequential division where micromeres cleave first, synchronously relative to each other, followed by asymmetric, synchronous cleavage of macromeres. Micromeres 1a and 1b cleave laeotropically and asymmetrically at a 45° angle, producing two progeny each: 1a^1^, 1a^2^, and 1b^1^, 1b^2^ respectively. The 1a^2^ and 1b^2^ blastomeres are smaller than 1a^1^ and 1b^1^. This results in an embryo with six cells – four micromeres and two macromeres. The 1A and 1B macromeres then cleave laeotropically and asymmetrically to produce micromeres 2a and 2b towards the animal pole and macromeres 2A and 2B on the vegetal pole, resulting in an 8-cell embryo. The 1a^1^, 1b^1^, 1a^2^ and 1b^2^ cells then cleave symmetrically and synchronously. The 1a^1^ and 1b^1^ cells cleave at a 45° angle to the AV axis, while 1a^2^ and 1b^2^ cleave perpendicular to the AV axis. These cleavages produce a 12-cell embryo. Next, a 14-cell embryo forms when the 2a and 2b cells cleave symmetrically. The macromeres then undergo an additional laeotropic, asymmetric division to produce the 3a and 3b micromeres and 3A and 3B macromeres, generating a 16-cell embryo. After the 16-cell stage, the “cap” of micromeres at the animal pole appears to move towards the vegetal pole as further cell divisions occur (17 - 22 hpl), enveloping the larger macromeres (Movie S1 and Movie S2). This envelopment of the macromeres represents the first instance of cell internalization during *Hofstenia* development. Individual blastomeres that result from the cell divisions after the 16-cell stage could not be discerned under a dissecting microscope.

### Morula stage (23-40 hpl)

Once the large macromeres are completely enveloped, the embryo becomes a spherical cluster of cells with macromeres occupying the interior; we refer to this as the Morula stage (Fig. 3A). This “ball” of cells is solid, with no blastocoel-like cavity detectable (Fig. S3A). The cells at the surface of the embryo display a rough, uneven texture. The majority of nuclei at this stage possess a distinct shape, with chromosomes arranged in a loose, circular pattern. The majority of nuclei at this stage possess a distinct shape, with diffuse chromosome appearing to be organized around a point, resembling a flower-like shape that we refer to as “rosettes” (Fig. S3B). Nuclei undergoing division, however, do not possess rosette-shaped nuclei, and instead have visible chromosomes, some appearing to be segregated to opposite poles, suggestive of anaphase.

### Dimple stage (41 - 55 hpl)

As cells continue to divide, a “dimple” forms on one side of the embryo, where cells appear to be internalized in time-lapse microscopy (Fig. 3A and Movie S1). Therefore, we refer to embryos during this time as the Dimple stage. This represents a second, previously undescribed cell internalization event during embryogenesis in an acoel. During the Dimple stage, a ring of concentrated actin forms at the site of the “dimple” (Fig. 3A and S2B). This ring of actin progressively becomes constricted as development continues during this stage, and as the “dimple” becomes smaller. Instead of the flowery, rosette-shaped nuclei that were detected in the Morula stage, many Dimple stage nuclei resemble a “donut”, forming a ring with an empty space at its center (Fig. S3B).

### Post Dimple stage (56 - 95 hpl)

The “dimple” becomes smaller in size, and the embryo acquires a smooth, spherical shape at the Post Dimple stage (Fig. 3A). Cells are now small enough to give the embryo’s surface an even, smooth texture. During this stage, the embryos do not exhibit gross morphological changes, while cell divisions continue. The nuclear rosettes/donuts are no longer detectable at this stage. The nuclei no longer have an empty space at their center, and adopt a more spherical shape (Figure S3B).

### Pill stage (96 - 104 hpl)

As cell division continues, the embryo becomes less spherical, adopting an oval shape, and can be observed spinning within its egg shell at the Pill stage (Fig. 3A, Movie S1). However, no body wall movement could be observed.

### Prehatchling stage (105 - 117 hpl)

The embryo continues to elongate, and the anterior-posterior axis is clearly defined by the presence of a mouth (anterior) and a tapered tail (posterior) at the Prehatchling stage (Fig. 3A). The embryo moves its body wall within its egg shell, suggesting the use of muscle (Movie S1). A region with sparsely-distributed nuclei is observed at the center of the embryo, which suggests the presence of the gut. A tube-like pattern of nuclei at the anterior of the embryo is also visible, which signifies the presence of a pharynx (Fig. 3A).

### Pigmented Prehatchling stage (118 - 192 hpl)

At the Pigmented Prehatchling stage, the embryo continues to elongate its body axis and the posterior becomes further tapered, creating the appearance of a Hatched Juvenile worm (Fig. 3A). Brown and white pigment granules form throughout the embryo. The embryo also moves vigorously within its egg shell. Furthermore, the space between the embryo and the egg shell diminishes, and the embryo could be seen pushed up against the internal wall of the shell just prior to hatching. The anterior region of the embryo is densely populated by nuclei at this stage, which corresponds to the anterior condensation of neurons that are present in the post-embryonic stages of *Hofstenia* (Hulett et al., 2020).

### Hatched Juvenile stage (193 - 204 hpl)

The embryo continues to increasingly occupy more space within the eggshell until it eventually breaks free as a Hatched Juvenile animal (Fig. 3A). There is variability in the timing of hatching, even among embryos from the same clutch, which are presumably fertilized at roughly the same time (Movie S1). Free-swimming Hatched Juvenile worms move on the substrate and swim through the water column. Although most of the worms acquire brown and white pigmentation, there is considerable phenotypic diversity in coloration (Fig. S1E). Some individuals possess little to no brown pigmentation, and appear mostly translucent with white bands. Others have very little white pigmentation, and are mostly brown. The white bands are not contiguous in some individuals, and are entirely absent in others.

#### Two distinct cell internalization events occur during Hofstenia development

Previous studies of acoel embryos reported a single cell internalization event where the vegetal macromeres are enveloped by the micromere progeny. This event is believed to represent gastrulation, as the internalized macromeres gave rise to the endomesoderm, and the conserved blastopore markers *brachyury* was detected at the site of internalization (Hejnol and Martindale, 2008a; Henry et al., 2000). Our staging series showed that *Hofstenia* embryos likely undergo two cell internalization events during early embryogenesis - 1) internalization of the vegetal macromeres at the Morula stage, and 2) internalization of cells at the Dimple stage (Fig. 3A and Movie S1). To facilitate better characterization of the two cellular internalization events, we utilized fluorescent dextran injections to visualize cells in developing embryos.

We injected the 1a^1^ and 1b^1^ micromeres of the 4-cell stage embryo with fluorescein dextran and imaged live the stages where cell movements were observed (Fig. 4). When viewed from the vegetal side during Early Cleavage (10hpl), the two unlabeled macromeres were seen as large, dark masses. As the labeled micromeres on the animal side continued to proliferate, their daughter cells were observed moving onto the vegetal side, and enveloping the two unlabeled macromeres (Fig. 4A and Movie S3). Thus, much like other acoel species, duet cleavage in *Hofstenia* concluded with the internalization of macromeres at the vegetal pole.

**Fig. 4.**
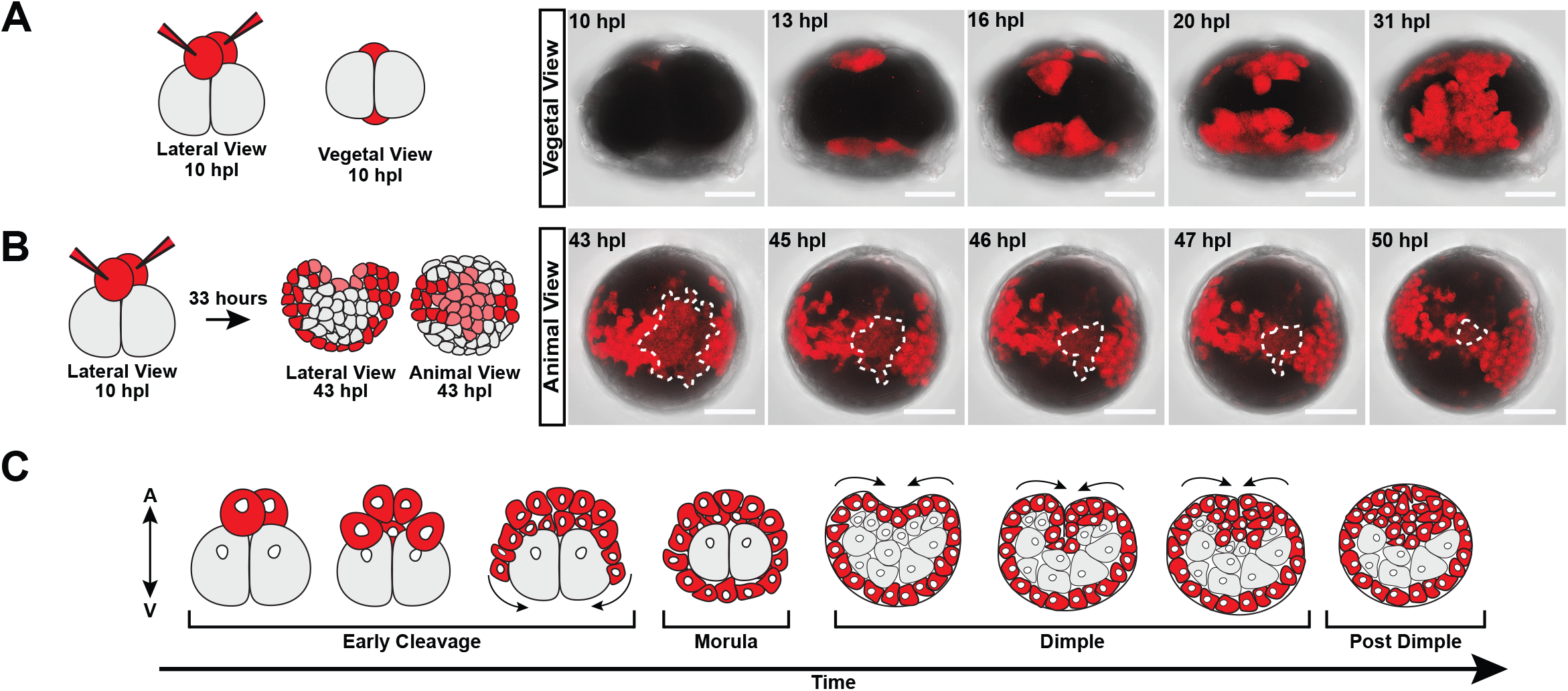
Two distinct cell internalization events occur during early development. (A) Left: Schematic depicting micromeres labeled at the 4-cell stage when viewed laterally and from the vegetal pole. Right: Timelapse screen captures of fluorescent dextran injected embryos when viewed from the vegetal pole (Movie S3). The daughter cells of micromeres labeled in red can be seen spilling over to the vegetal side and enveloping the macromeres (n=7). (B) Left: Schematic demonstrating the distribution of labeled cells at the Dimple stage when viewed laterally and from the animal pole. Right: Timelapse screen captures of dye injected embryos when viewed from the animal pole during the Dimple stage (Movie S4). The patch of cells outlined with a dashed white line can be observed being internalized (n=7). (C) Schematic model depicting the two cell internalization events during *Hofstenia* development, with all micromeres and their progeny colored in red. Scale bars, 100μm.

Next, we imaged the same micromere-injected embryos at the Dimple stage where a subset of labeled cells were observed to be internalized at the site of the “dimple” (Fig. 4B and Movie S4). This confirmed our initial observations that *Hofstenia* does indeed undergo a second, previously undescribed cell internalization event among acoels. The internalizing cells appeared to be less bright, retaining less of the fluorescent label relative to their neighboring cells that don’t become internalized. This suggests that the internalizing cells had undergone more cell divisions. Next, we performed continuous live-imaging of embryos (n=5) from Early Cleavage to the Dimple stage to determine where the second internalization, or the “dimple” occured in relation to the animal-vegetal axis of the embryo (Movie S5). We found that the “dimple” forms opposite to the site of macromere internalization (the first cell internalization). Therefore, this second cell internalization event occurs on the animal pole (Fig. 4C). Further studies of fate mapping in *Hofstenia* embryos are needed to determine whether either of these internalization events correspond to gastrulation in this species.

#### RNA sequencing suggests major transcriptional shifts occur after the Dimple stage

We next performed bulk RNA sequencing (RNA-seq) in triplicate on embryos spanning all major developmental stages to identify molecular correlates of the stages we defined based on morphology. All Early Cleavage stages were pooled together, and the Post Dimple stage was split into two different time ranges to gain further temporal resolution in gene expression. Principal component analysis (PCA) of our samples showed that replicates within the same developmental stages were consistent, enabling us to identify key differences across stages (Fig. 5A). Based on the spatial segregation of the two earliest stages sampled (Early Cleavage and Morula) on the PCA plot, we hypothesized that globally, the transcriptomes of these stages were distinct from those of all other samples. This suggested that a major transcriptional shift occurs at or preceding the Dimple stage, and we sought to identify the associated genes and molecular functions.

**Fig. 5.**
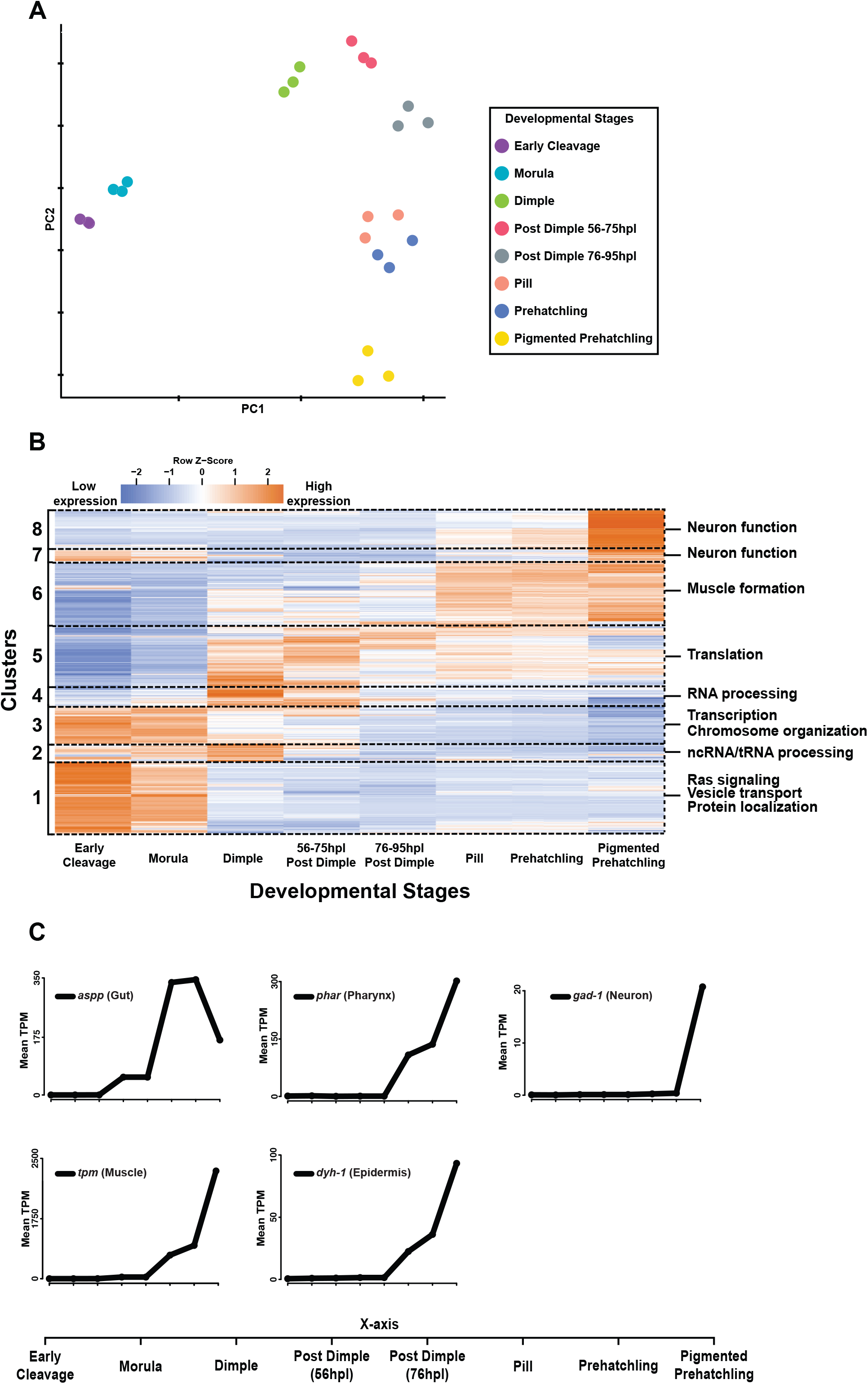
Bulk RNA-seq reveals suggests the formation of differentiated tissues occurs only after the Dimple stage. (A) PCA plot of all samples shows that the Early Cleavage and Morula stages are transcriptionally distinct from all other stages. Colors represent samples from the same developmental stage. (B) Heatmap of the mean transcripts per million (TPM) values across three biological replicates per developmental stage, of genes that are significantly differentially expressed during development (Likelihood Ratio Test). Columns represent developmental stages, rows represent genes. The dotted lines demarcate gene clusters generated from hierarchical clustering that have similar expression profiles across development. Each cluster identity is labeled on the left of the heatmap. Representative GO terms associated with each cluster are listed on the right of the heatmap. (C) Line graphs depicting mean transcript per million (TPM) values for known differentiated tissue marker genes. All markers shown have expression levels increasing only after the Dimple stage. The legend for the x-axis for all graphs is shown at the bottom.

To identify key genes that may have functional significance during *Hofstenia* development, we performed pairwise differential expression analyses between consecutive developmental stages and turned our attention to genes that changed significantly in expression during at least one transition during development. We then generated a heatmap plotting normalized expression values (transcripts per million, TPM) of these genes to visualize and identify broad patterns in expression during embryogenesis (Fig. 5B). Much like the PCA plot, the heatmap highlighted that the Early Cleavage and Morula stages were similar in gene expression profile, and were distinct from all other stages. The heatmap also revealed groups of genes that appeared to have similar expression dynamics across development, with hierarchical clustering recovering eight clusters with distinct patterns of expression.

We next sought to determine whether genes associated with specific functional roles could be enriched in these gene clusters. Thus, we utilized gene ontology (GO) enrichment analysis to capture any associated molecular functions (Fig. 5B and Table S2A). Notably, we found that clusters containing genes that became highly expressed starting at the Dimple stage were enriched for terms associated with translation, RNA processing, and muscle formation. Further, terms associated with neuronal function were enriched in clusters where the majority of the genes appeared to be highly expressed only late in development, at the Pigmented Prehatchling stage. The enrichment of terms associated with neuronal function and muscle formation among genes with elevated expression after the Dimple stage suggested that differentiated cell types were only present after this stage.

To confirm the GO enrichment analysis results and to expand on the hypothesis that differentiated cell types are only present after the Dimple stage, we examined the mean transcripts per million (TPM) values of known differentiated cell type markers across development (Fig. 5C). We found that all markers examined (gut: *aspp*, neuron: *gad-1*, epidermis: *dyh-1*, pharynx: *phar*, and muscle: *tpm*) increased in expression levels only after the Dimple stage. Furthermore, the expression profile of the muscle marker *tpm* and the neural marker *gad-1* mirrors the expression profile of clusters 6 and 8 which were found to be enriched for terms associated with muscle formation and neuronal function respectively. This suggests that organogenesis and substantial gene expression changes in the embryo occur after the Dimple stage, further highlighting the potential importance of the associated internalization event.

#### Cells expressing differentiated cell markers appear after the Dimple stage

We performed *in situ* hybridization of the differentiated tissue markers examined above to assess the hypothesis that these markers are expressed only after the Dimple stage, and to determine where within the embryo these markers were expressed. We found that the expression of gut, muscle, epidermis, pharynx, and neural markers were all detected after the Dimple stage (Fig. 6A,B).

**Fig. 6.**
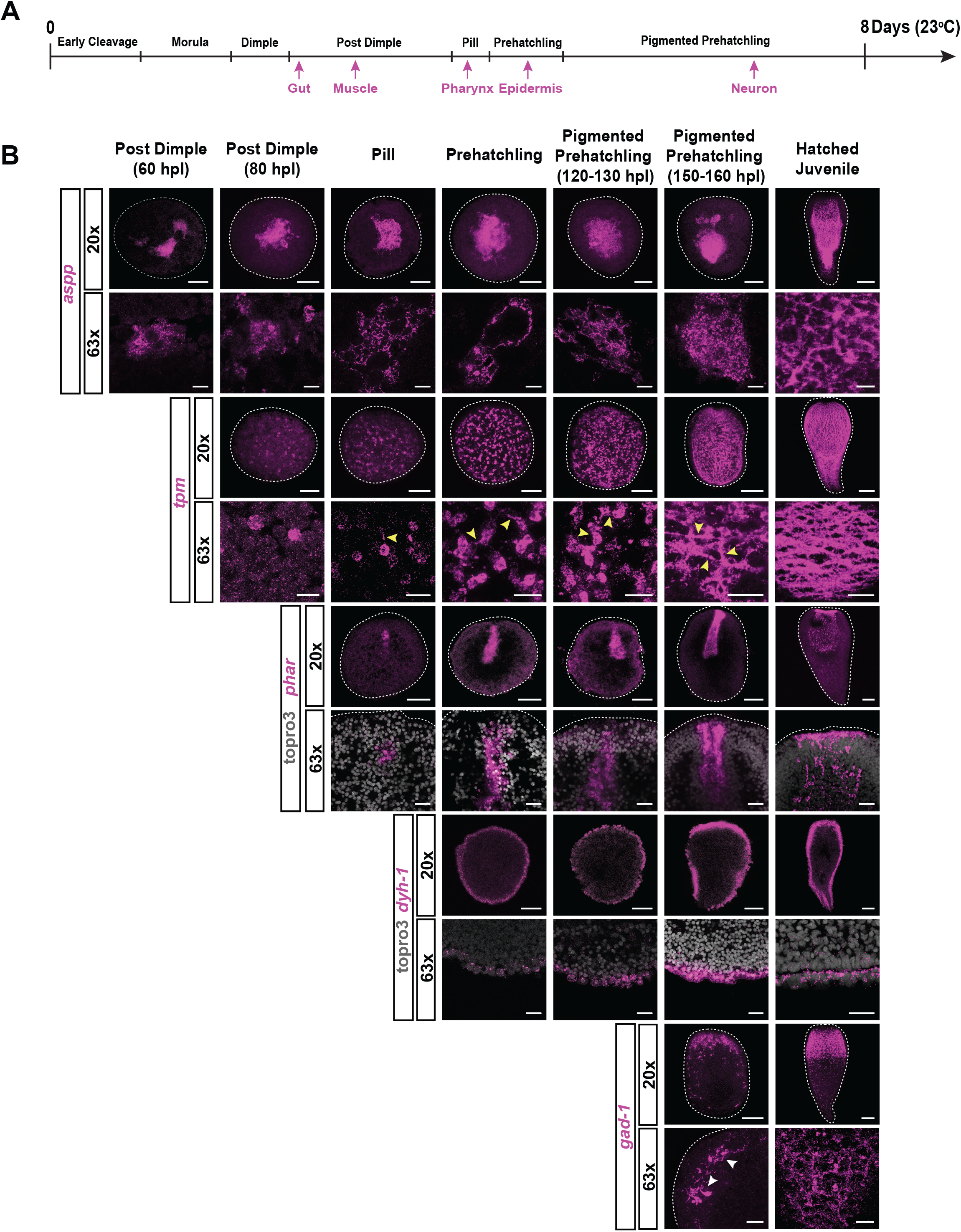
Expression of differentiated tissue marker genes was detected after the Dimple stage. (A) Schematic timeline of development showing when markers for differentiated cell types were first detected via *in situ* hybridization. (B) *in situ* hybridization of differentiated cell markers. *aspp* marked the gut in the Hatched Juvenile worms (z-projection), and internal cells in embryos (optical section) starting from the Post Dimple stage (60hpl). *tropomyosin* marked muscle in the Hatched Juvenile worms (z-projection), showing a mesh-like network of muscle fibers. Expression was first detected at the Post Dimple stage (80hpl), and cells started to exhibit fiber-like projections at the Pill stage (yellow arrows) (optical section). The pharynx marker *phar* was first detected at the Pill stage as a patch of cells on one side of the embryo (optical section). This patch of expression changed into a tube-like structure that extended internally at the Prehatchling stage, outlining the pharynx. The nuclear stain topro3 was used in 63x magnification images. *dyh-1* marked the epidermis in the Hatched Juvenile (optical section), and expression was detected in the outermost cells of the embryos at the Prehatchling stage (optical section). Topro3 was used in the 63x magnification images to visualize the outer boundaries of the embryo. *gad-1* marked neural cell types in the Hatched Juvenile (z-projection), showing a high condensation of cells in the anterior compared to a more diffuse pattern of expression throughout the rest of the body. *gad-1* was only detected at 150hpl Pigmented Prehatchling (z-projection), showing expression patterns resembling that of the Hatched Juveniles, and axon-like extensions being visible (white arrows). Scale bars, 100μm (20x magnification) and 25 μm (63x magnification). Dashed-lines represent the outer boundary of embryos and Hatched Juveniles.

In the Hatched Juvenile stage *Hofstenia*, *aspp* (*lysosomal aspartic protease*) marked the gut, revealing cells that were closely associated with one another in the internal region of the worm (Fig. 6B). In embryos, *aspp* expression was detected among a tight cluster of cells that occupied the center of the embryo starting at the early Post Dimple stage (60hpl). Imaging at higher magnification showed an increase in the abundance of these cells with time.

The muscle marker *tpm* (*tropomyosin*) marked a mesh-like network of orthogonal muscle fibers that were present in the sub-epidermal periphery and pharynx of the Hatched Juvenile stage worm (Fig. 6B, Fig. S4). Among embryos, *tpm* also occupied the sub-surface periphery, and was first detected at the late Post Dimple stage (80 hpl) (Fig. 6B, Fig. S4). High magnification imaging revealed the gradual emergence of cellular extensions starting at the Pill stage, which ultimately formed a mesh-like network resembling that of the Hatched Juvenile stage at the late Pigmented Prehatchling stage (150 - 160 hpl). This suggests that although cells that express *tpm* are present as early as the Post Dimple stage (80 hpl), they do not necessarily adopt the morphology of mature muscle tissue until very late in development.

The *Hofstenia* specific gene *phar* (*pharynx*) clearly labeled both the opening and internal structure of the pharynx at the Hatched Juvenile stage (Fig. 6B). This gene was detected at the Pill stage as a patch of cells on the presumptive anterior of the embryo. As the embryo developed further, *phar* was observed in a tube-like pattern which extended internally from the outer surface of the embryo (Fig. 6B). The timing of *phar* expression in a pharynx-like pattern matched with when we were able to detect the formation of the mouth in our live-imaging and developmental atlas (Fig. 3A, Movie S1).

The epidermal marker *dyh-1* (*dynein heavy chain-1*) was expressed in the epidermis of the Hatched Juvenile worm (Fig. 6B). Expression was detected on the outer periphery of the embryo from the Prehatchling stage. High magnification images confirmed that *dyh*-*1* was only expressed in the outermost cell layer and this expression was maintained until hatching. This suggested that *dyh-1* was indeed marking the cells that ultimately became the mature epidermis. Finally, the neural marker *gad-1* (glutamate decarboxylase) marks the anterior condensation and body neurons in hatched worms, as described by (Hulett et al., 2020). During development, this gene was only detectable at the late Pigmented Prehatchling stage (150 - 160 hpl), with the distribution of signal resembling that of a Hatched Juvenile stage. Higher magnification images confirmed the presence of axon-like extensions in both the Pigmented Prehatchling stage and Hatched Juvenile stages.

The timing or stage of detection of these markers through *in situ* hybridization corroborated the temporal dynamics of the expression levels of these genes as revealed by the RNA-seq data shown above (Fig. 5C and Fig. 7). Additionally, the markers that were detected correspond to tissues derived from all three germ layers that are present among bilaterian animals. This could suggest that the specification of the three germ layers occurs after the Dimple stage. However, we cannot rule out the possibility that germ layers are specified earlier, possibly even before the Dimple stage, with differentiated cell type markers becoming expressed only later in development. Once the markers of differentiated cell types were observable, we noted substantial changes in these tissues. For example, gut, muscle, and pharyngeal cells became more numerous and organized into distinct structures (Fig. 6B). Based on these data, we infer that organogenesis occurs after the Dimple stage.

**Fig. 7.**
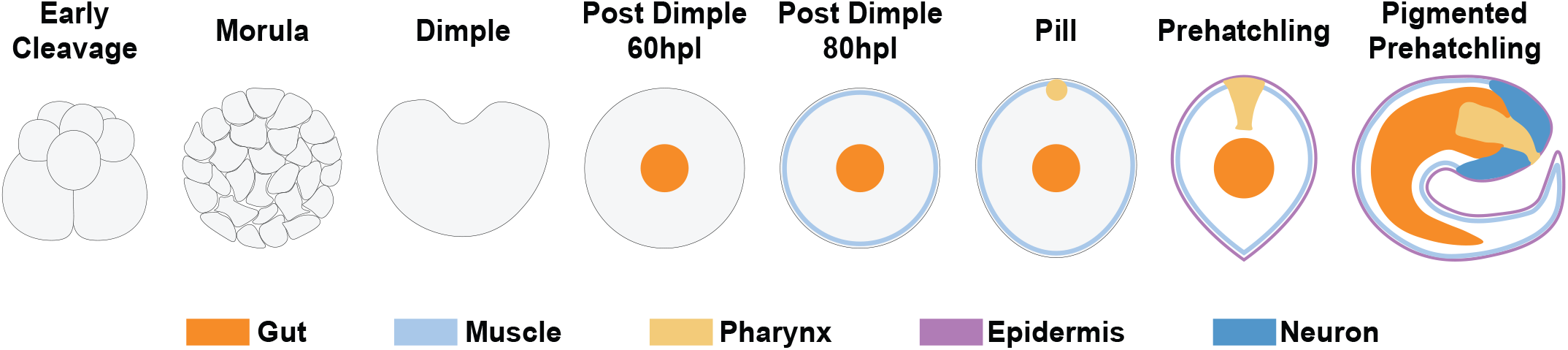
Schematic summarizing major developmental stages and formation of differentiated tissues. During *Hofstenia* development the gut was first detected in the center of the embryo (orange). Next, muscle was detected in the sub-surface periphery of the embryo (light blue). The pharyngeal marker was then detected in a small patch on the surface of the embryo (yellow). The epidermal marker was detected at the Pill stage (purple). Finally, neurons were detected at the Pigmented Prehatchling stage internally on the anterior side of the embryo (dark blue).

## Discussion

This study provides an in-depth characterization of embryogenesis of the acoel worm *Hofstenia miamia*. *Hofstenia* embryos undergo duet cleavage, a stereotyped form of early embryonic development that is specific to acoels. Although all duet cleavage programs involve the formation of a “duet” of animal pole micromeres and a pair of vegetal pole macromeres, acoel species display variations in the number, order, and symmetries of cleavages. We found that *Hofstenia* embryos cleave in a stereotyped pattern that differs slightly from the patterns reported from other species. Previous studies of duet cleavage in other species have shown, in some instances, the stereotyped formation of strings of macromeres or blastomere fusion events in an otocelid acoel (*Archocelis macrorhabditis*) and an isodiametrid acoel (*Archaphanostoma agile*) respectively (Apelt, 1969). *Hofstenia* exhibited fusion events on occasion, but these events were not a part of the stereotyped cleavage program. The *Hofstenia* cleavage pattern was most similar to those of the convolutid species *Neochildia fusca*, *Convolutriloba longifissura*, and *Symsagittifera roscoffensis*, but showed some differences (Bresslau, 1909; Hejnol and Martindale, 2008a; Henry et al., 2000). At the four-cell stage in *Hofstenia*, the micromeres 1a and 1b cleave before the 1A and 1B macromeres, whereas the macromeres cleave before the micromeres in convolutids. Moreover, the cleavage of the 1a and 1b micromeres in *Hofstenia* was asymmetric, in contrast to the symmetric cleavage of the corresponding micromeres observed among the convolutids. Furthermore, macromeres 3A and 3B are internalized at the 16-cell stage in convolutids. In *Hofstenia,* the macromeres were not fully internalized until after subsequent cell divisions. Internalization of the vegetal macromeres in *Hofstenia* embryos started only after the 16-cell stage was reached (Movie S2). Given that *Hofstenia* represents a sister-group to the vast majority of acoels whereas convolutids diverge later within the clade (Jondelius et al., 2011), we infer that the conserved features of duet cleavage in acoels include the formation of a pair of vegetal macromeres that cleave counter-clockwise and asymmetrically to form “duets’’ of smaller micromeres on the animal side which ultimately envelop the vegetal macromeres. Fate mapping studies in multiple species are needed to assess if the shared macromere-micromere organization of the embryos represents conserved specification events across acoels.

The duet cleavage pattern observed in *Hofstenia* and other acoel embryos is reminiscent of spiral cleavage, which involves the formation of four vegetal macromeres that produce “quartets” of micromeres toward the animal pole. The former placement of acoels within the spiralian phylum Platyhelminthes and the striking similarities between spiral and duet cleavage suggested that duet cleavage is a derived form of spiral (Ax and Dörjes, 1966; Ax and Jeffries, 1987; Boyer et al., 1996; Boyer and Jonathan, 1998; Bresslau, 1909; Costello and Henley, 1976; Henry and Martindale, 1999; Hyman, n.d.; Peterson and Eernisse, 2001; Smith et al., 1986). With molecular phylogenies now consistently placing acoels as distantly related to Platyhelminthes, duet cleavage should not be studied with an assumption of homology to spiral cleavage (Hejnol et al., 2009; Jondelius et al., 2011; Kapli and Telford, 2020; Marlétaz et al., 2019; Mwinyi et al., 2010; Philippe et al., 2007, 2011, 2019; Ruiz-Trillo et al., 1999, 2002, 2004; Ruiz-Trillo and Paps, 2016; Sempere et al., 2007; Telford et al., 2003). However, given the similarities in cleavage programs between these distantly related lineages, detailed comparisons of the cell and molecular mechanisms underlying spiral and duet cleavage could uncover conserved mechanisms that represent ancestral states of bilaterian development. *Hofstenia* embryos, which are amenable to experimental manipulations, will facilitate systematic comparisons of duet and spiral cleavage programs.

Our developmental atlas showed that after duet cleavage is completed, *Hofstenia* undergoes two cell internalization events. The first of these cell internalization events occurred through the movement of animal pole micromeres which resulted in the vegetal macromeres being internalized. The second internalization event, which had not been described previously in other acoel species, occurred at the Dimple stage at the animal pole. Given the two cell internalization events observed during *Hofstenia* development, the event corresponding to the formation of three germ layers, *i.e.*, gastrulation, remains to be identified. Previous work defined gastrulation in acoels as the internalization of the macromeres, a direct parallel to the mechanism of gastrulation in some spiralian embryos (Apelt, 1969; Bresslau, 1909; Hejnol and Martindale, 2008a; Henry et al., 2000). However, spiralians undergo gastrulation in different modes, despite their conserved cleavage program (*e.g.* epiboly and invagination at different stages of development) (Lambert, 2010; Lyons and Henry, 2014). It is possible that even among acoels, the mode of gastrulation differs widely. It remains unknown whether this second internalization has remained undetected in previously studied species, or whether *Hofstenia* represents a unique instance where a second internalization event occurs at the Dimple stage. Our work shows that this cellular event corresponds to a major transcriptional shift where the gene expression profile becomes distinct from that of the earlier Early Cleavage and Morula stages. Moreover, GO enrichment analysis and *in situ* hybridization data detected the expression of differentiated tissue marker genes corresponding to all three germ layers only after the Dimple stage (Fig. 5C; Fig. 6; and Fig. 7). Further work on fate-mapping of cells internalized at the Dimple stage as well as studies of the function of conserved genes associated with gastrulation is needed to understand how *Hofstenia* gastrulates.

Our *in situ* hybridization data showed when differentiated tissue types were present during *Hofstenia* embryogenesis. Previous studies on other acoel species have highlighted the formation of muscle fibers and the expression of neural transcription factors during embryonic development. We found that the pattern by which *Hofstenia* muscle cell extensions form differs from the process of muscle formation in other acoels such as *S. roscoffensis* and *C. pulchra*. In *C. pulchra*, circular, latitudinal muscle fibers appear first, and are later connected by longitudinal fibers (Ladurner and Rieger, 2000). Myogenesis in *S. roscoffensis* occurs by the formation of muscle fibers sequentially from the animal to the posterior pole (Semmler et al., 2008). We did not observe sequential formation of muscle in *Hofstenia*. Instead, the fibers were first distributed in random directions and across all parts of the embryo, and then resolved into an organized lattice by 150 hpl. Formation of neurons has been inferred via studies of the expression of basic helix-loop-helix (BHLH) transcription factors, often associated with neurogenesis, in the embryos of the acoel *S. roscoffensis (Perea-Atienza et al., 2018)*. In *S. roscoffensis*, the majority of the BHLH transcription factors tested were expressed within the first 24 hours of development. This represents expression at an early stage, given that it takes 4-5 days for this species to complete embryogenesis. This study most likely found early progenitors of neurons, while our study focused on identifying when mature neurons were present. Further studies of neural transcription factor expression in *Hofstenia* embryos are needed to determine when neural progenitors are present in this species. Although neuron and muscle formation in other acoel embryos have been studied as described above, the timing for the formation of other differentiated cell types/organs such as gut, epidermis, and pharynx have not been investigated before.

*Hofstenia* represents a genomically-enabled, early-diverging member of the acoel clade, making its embryos an attractive system for functional studies that inform the evolution of development (Gehrke et al., 2019; Jondelius et al., 2011; Srivastava et al., 2014). Further, given *Hofstenia*’s capacity for whole-body regeneration, its embryos can address questions of stem cell specification, maintenance, and differentiation. This work represents the first step in establishing *Hofstenia* embryos as a new acoel model for developmental biology.

## Materials and Methods

### Hofstenia miamia adult and embryo culture

*Hofstenia miamia* adults were cultured in plastic boxes in artificial sea water at 21°C, and embryos were found typically laid on the plastic in clutches (Fig. S1B,C). Embryos were collected using a glass Pasteur pipette to scrape them off of surfaces. Embryos at the 2- and 4-cell stages were sorted manually under a dissection microscope, and incubated at 23°C until the desired stage was reached. Embryos were staged based on morphological changes observed under a dissection microscope and developmental timing determined based on the number of hours post laying at 23°C.

### Embryo de-shelling and fixation

In order to allow penetration of *in situ* probes and small molecules past the egg shell, embryos were treated with a de-shelling solution (32mM sodium hydroxide, 0.5mg/ml sodium thioglycolate, and 1mg/ml of pronase in artificial seawater) for varying times depending on developmental stage while being placed on a shaker. Embryos from Early Cleavage to Pill stages were incubated for eight minutes, while Prehatchling and Pigmented Prehatchling stages were incubated for six minutes. Once treated, embryos were fixed in 4% paraformaldehyde solution in seawater overnight at 4°C. After fixation, embryos were stored in phosphate buffered saline (PBS) in the 4°C fridge for a maximum of one week before use in *in situ* hybridization.

### Dye injections

A plastic mold with 300μm pins was made using a laser cutter (Ricci et al., 2020, *in prep*). This mold was placed in molten 1% agarose. Once the agarose solidified, the pins created small holes into which embryos were placed. Embryos were then injected with fluorescein dextran using a Narishige micromanipulator connected to a Harvard Apparatus injector and quartz needles (Sutter Instrument GF100-50-10).

### Live imaging

Embryos were mounted on glass slides underneath coverslips with “clay feet”. The slides were then sealed using vaseline. Embryos that were injected with fluorescein dextran were imaged with a Leica SP8 confocal microscope at intervals of 10 minutes. For (Movie S1), (Movie S2), and (Movie S5), images were taken at intervals of 2, 5, and 10 minutes respectively on a Leica DM8000 stereomicroscope.

### RNA sequencing

RNA was extracted from eight stages (Early Cleavage, Morula, Dimple, Post Dimple 56-75hpl, Post Dimple 76-95hpl, Pill, Prehatchling, and Pigmented Prehatchling) using the Nucleospin RNA XS kit (Macherey Nagel). To obtain enough total RNA to synthesize libraries, multiple embryos (approximately 90-100) were pooled together to generate a single biological replicate for each stage. Three biological replicates were made for each developmental stage. RNA-seq libraries were generated using the Illumina TruSeq-v2 kit (RS-122-2001 and RS-122-2002). Libraries were analyzed for quality using the Agilent Tapestation High Sensitivity D1000 tape, and were subsequently used for single-end, 75bp sequencing in one lane of the Illumina NextSeq sequencer.

Libraries were de-multiplexed and reads were pseudo-mapped and quantified using Salmon (Patro et al., 2017). Salmon outputs were then converted into a format that can be used by Sleuth using the R package Wasabi. The Wasabi R package can be found here: https://github.com/COMBINE-lab/wasabi. Differential expression was called with the R package Sleuth using the likelihood ratio test (Pimentel et al., 2017). Differential expression analysis was performed in a sequential fashion, performing pairwise comparisons of developmental stages adjacent in time to one another. Differential expression was called by setting a qvalue cutoff of 0.05, with the exception of the comparison between Pill and Prehatchling stages, where a pvalue cutoff of 0.05 was used (Table S2B). All hierarchical clustering and heatmap generation was performed using the gplots package in R. The heatmap was generated using a matrix of TPM values of genes with at least one statistically significant difference when comparing consecutive stages during development (Table S2C). Hierarchical clustering was done using the WardD clustering method on this matrix using the hclust function. Principal component analysis was performed on the sleuth object generated using estimated counts. Generation of line graphs depicting mean TPM values across three biological replicates were done in R using a custom script. All mean TPM values were calculated by taking the mean of the TPM values in each biological replicate for each developmental stage (Table S2D). GO enrichment analysis was performed using DAVID functional annotation tool (Table S2A). The best BLAST hit for *Hofstenia* genes against humans were used to perform GO enrichment analysis. The *Hofstenia* transcriptome was used as the background in this analysis. R scripts that have been used to call differential expression, generate plots, and perform hierarchical clustering can be found on github: https://github.com/JulianKimura/R_scripts

### Gene cloning

All genes were annotated based on the best BLAST hit to humans and were given UniProt identifiers. Genes of interest were amplified and cloned using the methods described in (Srivastava et al., 2014). Gene were named using BLAST-based sequence similarity to known proteins, and their corresponding primer and sequence information are listed in (Table S1).

### In situ hybridization

Riboprobes used for *in situ* hybridization were synthesized using the methods described by (Pearson et al., 2009). An adjusted version of the *in situ* hybridization protocol mentioned in (Srivastava et al., 2014) was used on de-shelled and fixed embryos. During the proteinase K step, embryos were treated for varying times depending on the stage, 1min for 0-107hpl, and 3mins for 108hpl-Hatched Juvenile. The pre-hybridization and hybridization solutions were made using 8M urea instead of DI formamide (Sinigaglia et al., 2018). With the exception of (Fig. S3A) which was stained with DAPI, all nuclei were stained using a TO-PRO™-3 Iodide (642/661) (T3605, Thermofisher).

### Phalloidin staining

Phalloidin staining was performed on de-shelled and fixed embryos using the methods described in (Srivastava et al., 2014).

## Acknowledgements

The optimization of microinjections was done with the help of Dr. Mark Martindale. We thank Dr. Seth Donoughe and Dr. Cassandra Extavour for helpful discussions, making the embryo microinjection molds, and inspiration for the developmental time series. We would also like to thank Dr. Andrew Gehrke, Dr. Marcela Bolaños, Dr. Yi Jyun Luo, Dr. Alyson Ramirez, Ryan Hulett, and Dr. Deirdre Lyons for helpful discussions. Finally, we would like to acknowledge the intellectual and technical support from all members of the Srivastava Lab.

## Data Availability

Raw fastq files of the embryonic transcriptome were submitted to the NCBI Sequence Read Archive under the bioproject code PRJNA603318. Gene sequences were submitted to genbank.

---

**Fig. S1.**
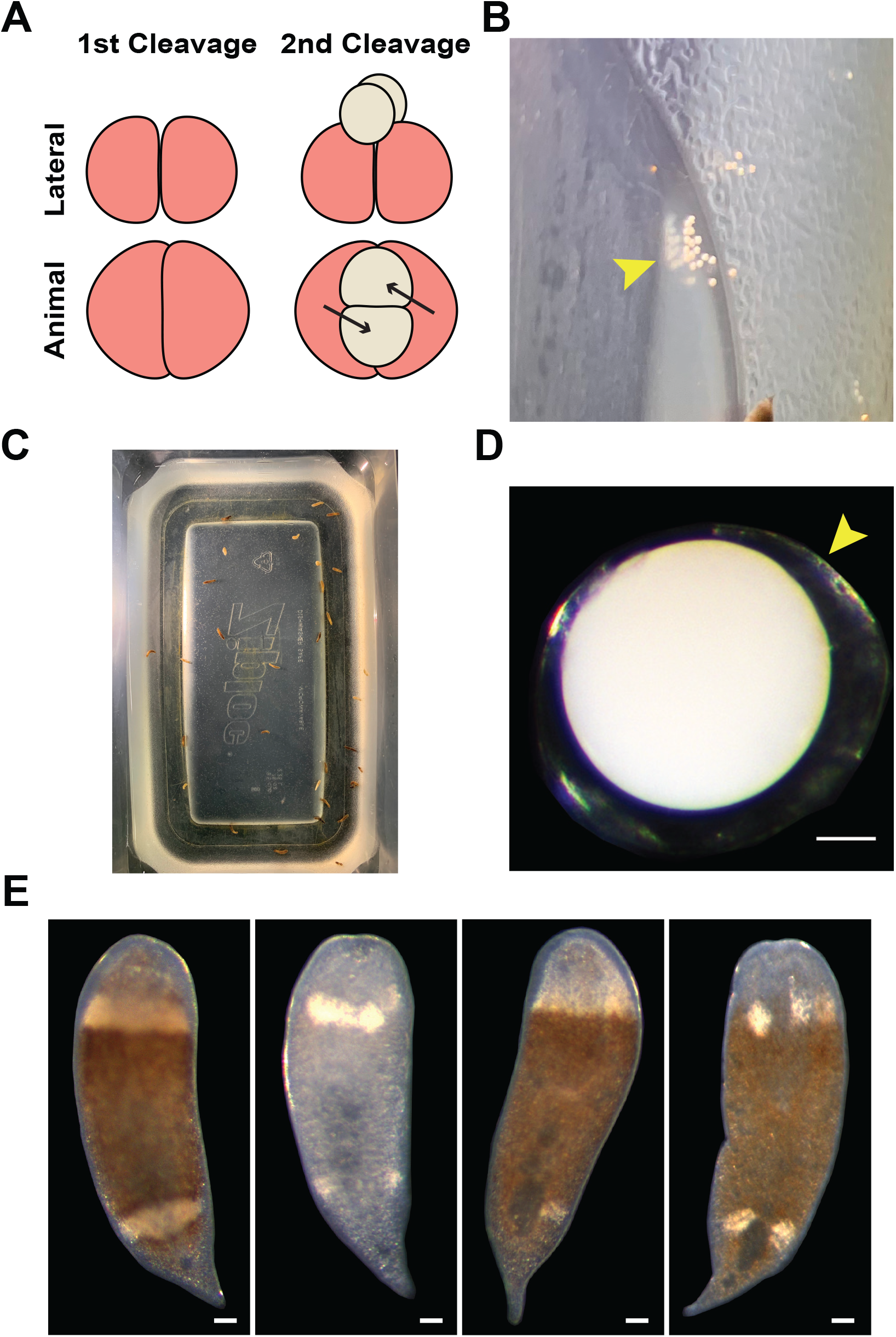
Culturing *Hofstenia* embryos. (A) Schematic illustrating the first two cleavages that occur during duet cleavage. Arrows denote the counter-clockwise direction of cleavage when viewed from the animal pole. The micromeres (yellow) are situated at the cell junction of the macromeres (red). (B) Clutch of embryos that were laid on the side of the tupperware box container. (C) Inside of a tupperware box used for culturing sexually mature *Hofstenia*. (D) Darkfield image of a zygote inside of a clear egg shell (yellow arrow). Scale bar, 100μm. (E) The phenotypic diversity in coloration that was detected among *Hofstenia* juveniles.

**Fig. S2.**
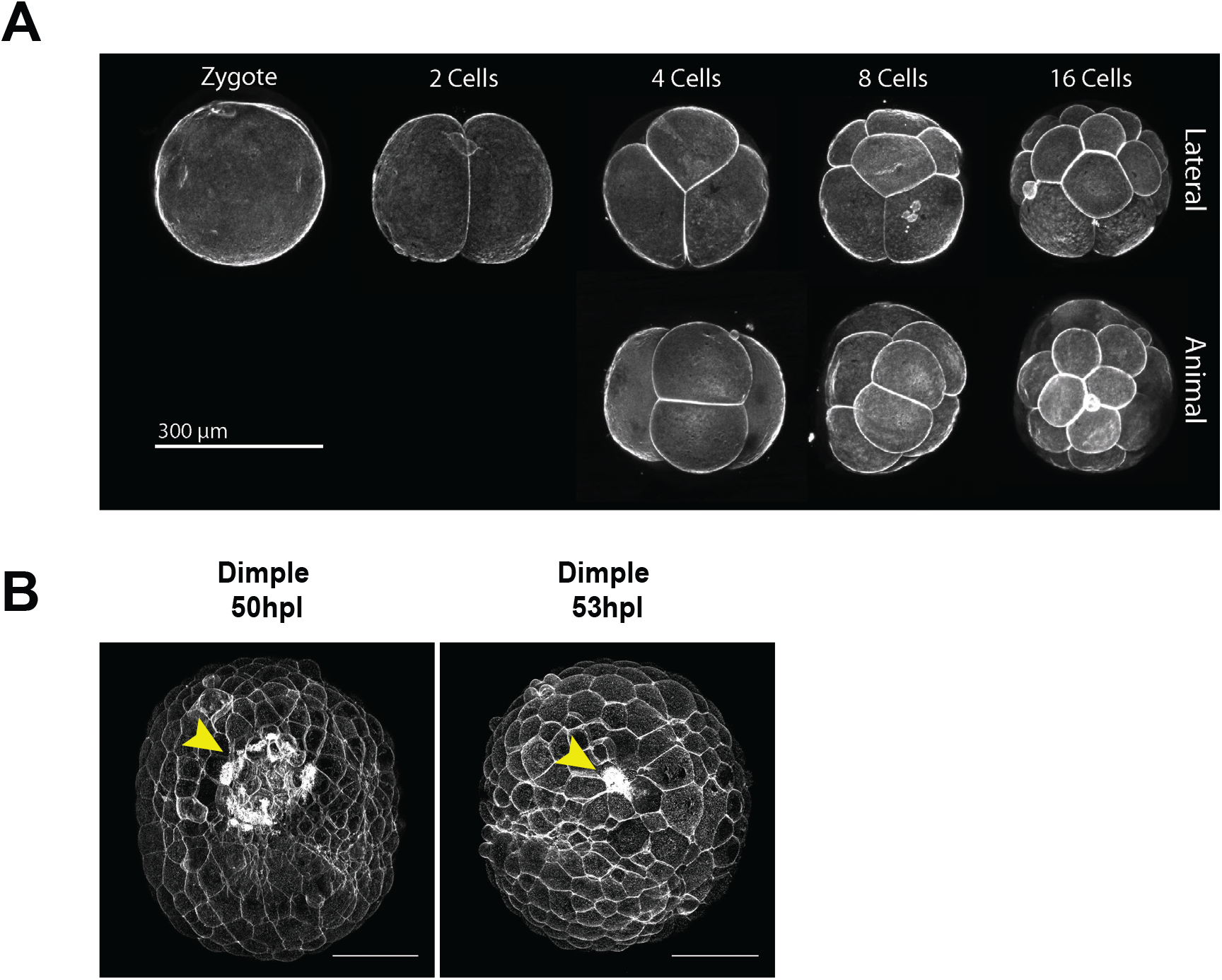
Phalloidin staining of *Hofstenia* embryos. (A) Representative images of phalloidin stained early cleavage embryos. Scale bar, 300μm (B) Phalloidin stained Dimple stage embryos viewed from the animal pole. A ring of actin is present at the site of cell internalization, or the “dimple”. This ring becomes progressively smaller as the embryo progresses through the Dimple stage, and ultimately disappears by the Post Dimple stage. Scale bars, 100μm

**Fig. S3.**
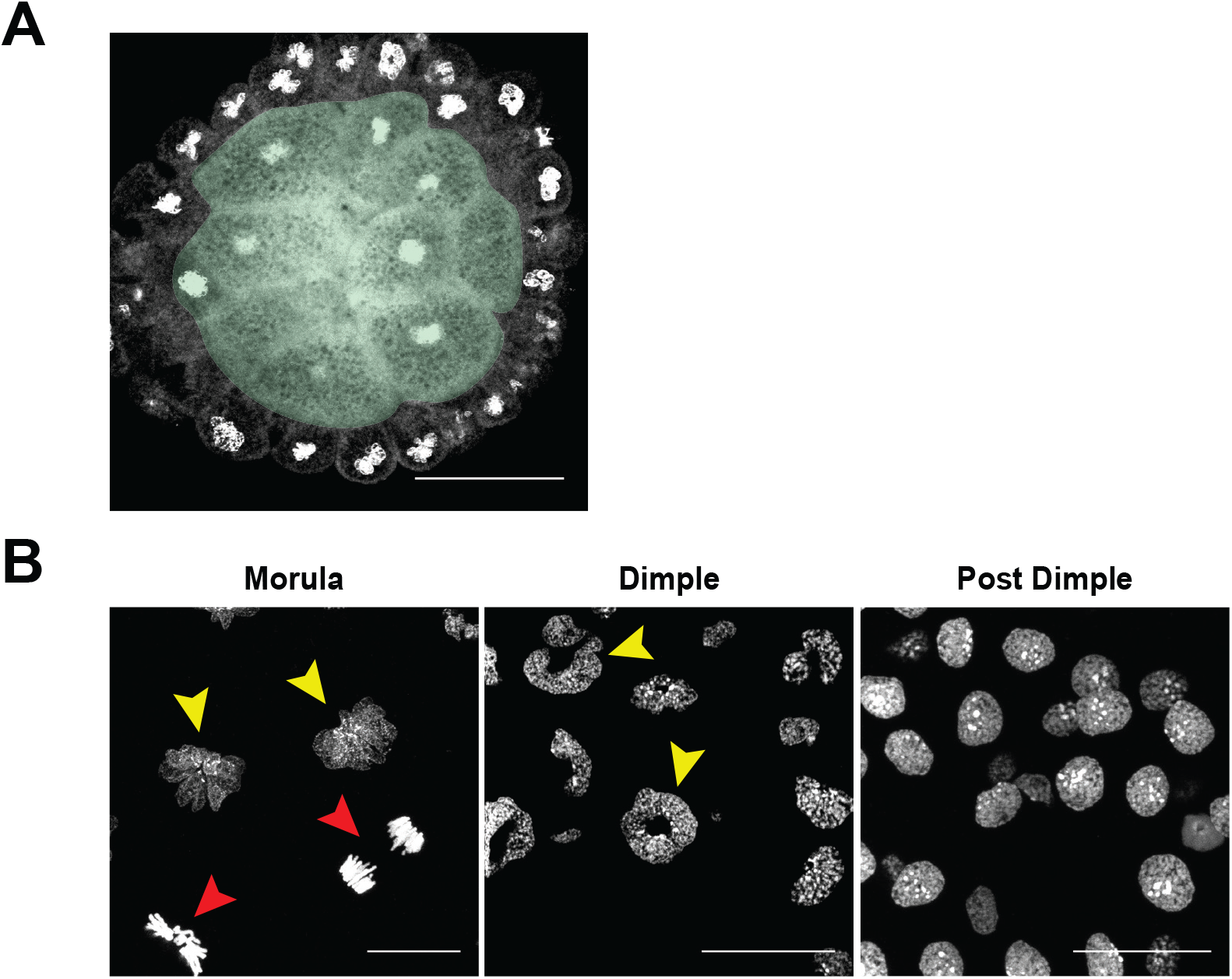
Nuclear staining of *Hofstenia* embryos. (A) A cross section of a DAPI stained Morula stage embryo shows the lack of a blastocoel-like cavity, with the internal space of the embryo being occupied by large cells (shaded green). (B) 63x imaging of Topro3 stained nuclei in Morula, Dimple, and Post Dimple stages. A rosette-like shape is present in the majority of Morula embryos (yellow arrow), with the exception of cells in anaphase (red arrow). Dimple stage embryos possess nuclei that are donut shaped (yellow arrow). The nuclei here are circular with an empty space at their centers. By the Post Dimple stage nuclei no longer have a rosette or donut shape, and resemble puncta. Scale bars, 25μm

**Fig. S4.**
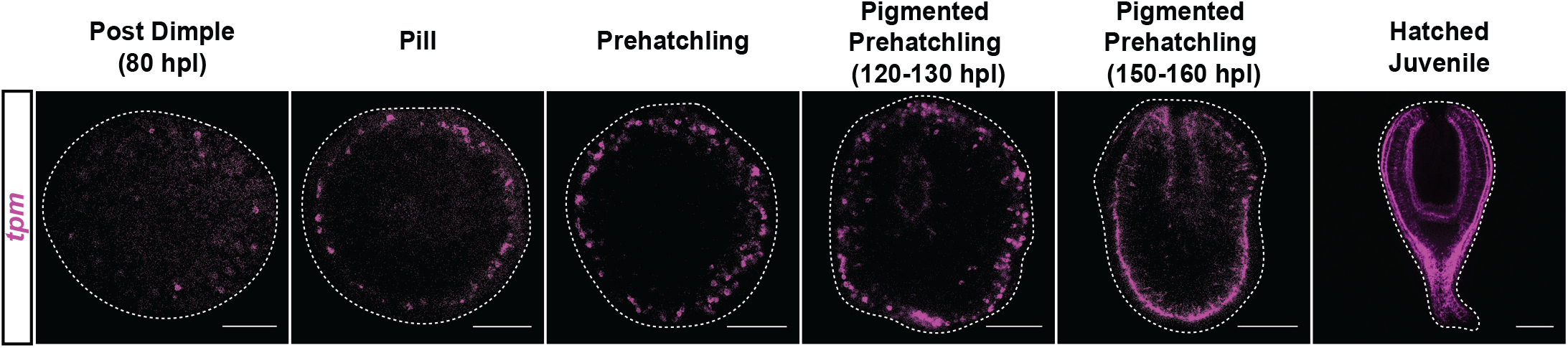
Cross sections of embryos showing *tropomyosin* expression *in situ*. Optical cross sections of *tpm in situ* hybridizations on embryos. Cells expressing *tpm* are situated below the surface of the embryo, but are distributed around the periphery of the cross section. This creates a ring of *tpm* expressing cells when viewed from a cross section. Scale bars, 100μm

## References

Apelt G (1969) Fortpflanzungsbiologie, Entwicklungszyklen und vergleichende Frühentwicklung acoeler Turbellarien. Marine biology 4(4). Springer-Verlag: 267–325.

Ax P and Dörjes J (1966) Oligochoerus limnophilus nov. spec., ein kaspisches Faunenelement als erster Süßwasservertreter der Turbellaria Acoela in Flüssen Mitteleuropas. Internationale Revue der gesamten Hydrobiologie und Hydrographie 51(1). Wiley: 15–44.

Ax P and Jeffries RPS (1987) The phylogenetic system: the systematization of organisms on the basis of their phylogenesis. Wiley New York. Available at: https://pdfs.semanticscholar.org/be84/045da27d1f14174891a91c9b53e643470dc6.pdf.

Bourlat SJ and Hejnol A (2009) Acoels. Current biology: CB 19(7): R279–80.

Boyer BC (1971) Regulative development in a spiralian embryo as shown by cell deletion experiments on the Acoel, Childia. The Journal of experimental zoology 176(1): 97–105.

Boyer BC and Jonathan QH (1998) Evolutionary modifications of the spiralian developmental program. American zoologist 38(4). Oxford University Press (OUP): 621–633.

Boyer BC, Henry JQ and Martindale MQ (1996) Modified Spiral Cleavage: The Duet Cleavage Pattern and Early Blastomer Fates in the Acoel Turbellarian Neochildia fusca. The Biological bulletin 191(2): 285–286.

Bresslau E (1909) Die Entwicklung der Acoelen. Deutsche Zoologische Gesellschaft 19: 314–324.

Costello DP and Henley C (1976) Spiralian Development: A Perspective. American zoologist 16(3). Oxford University Press (OUP): 277–291.

Gehrke AR and Srivastava M (2016) Neoblasts and the evolution of whole-body regeneration. Current opinion in genetics & development 40: 131–137.

Gehrke AR, Neverett E, Luo Y-J, et al. (2019) Acoel genome reveals the regulatory landscape of whole-body regeneration. Science 363(6432). DOI: 10.1126/science.aau6173.

Hejnol A and Martindale MQ (2008a) Acoel development indicates the independent evolution of the bilaterian mouth and anus. Nature 456(7220): 382–386.

Hejnol A and Martindale MQ (2008b) Acoel development supports a simple planula-like urbilaterian. Philosophical transactions of the Royal Society of London. Series B, Biological sciences 363(1496): 1493–1501.

Hejnol A and Pang K (2016) Xenacoelomorpha’s significance for understanding bilaterian evolution. Current opinion in genetics & development 39: 48–54.

Hejnol A, Obst M, Stamatakis A, et al. (2009) Assessing the root of bilaterian animals with scalable phylogenomic methods. Proceedings of the Royal Society B: Biological Sciences. DOI: 10.1098/rspb.2009.0896.

Henry JJ and Martindale MQ (1999) Conservation and innovation in spiralian development. Hydrobiologia 402(0): 255–265.

Henry JQ, Martindale MQ and Boyer BC (2000) The unique developmental program of the acoel flatworm, Neochildia fusca. Developmental biology 220(2): 285–295.

Hulett RE, Potter D and Srivastava M (2020) Neural architecture and regeneration in the acoel Hofstenia miamia. Proceedings. Biological sciences / The Royal Society 287(1931): 20201198.

Hyman L (n.d.) The Invertebrates, Vol II: Platyhelminthes and Rhynchocoela. 1951. New York: McGraw-Hill Book Company, Inc.

Jondelius U, Wallberg A, Hooge M, et al. (2011) How the worm got its pharynx: phylogeny, classification and Bayesian assessment of character evolution in Acoela. Systematic biology 60(6): 845–871.

Kapli P and Telford MJ (2020) Topology-dependent asymmetry in systematic errors affects phylogenetic placement of Ctenophora and Xenacoelomorpha. Science advances 6(50). DOI: 10.1126/sciadv.abc5162.

Ladurner P and Rieger R (2000) Embryonic Muscle Development of Convoluta pulchra (Turbellaria–Acoelomorpha, Platyhelminthes). Developmental biology 222(2): 359–375.

Lambert JD (2010) Developmental patterns in spiralian embryos. Current biology: CB 20(2): R72–7.

Lyons DC and Henry JQ (2014) Ins and outs of Spiralian gastrulation. The International journal of developmental biology 58(6–8): 413–428.

Marlétaz F, Peijnenburg KTCA, Goto T, et al. (2019) A New Spiralian Phylogeny Places the Enigmatic Arrow Worms among Gnathiferans. Current biology: CB 29(2): 312–318.e3.

Maslakova SA, Martindale MQ and Norenburg JL (2004) Vestigial prototroch in a basal nemertean, Carinoma tremaphoros (Nemertea; Palaeonemertea). Evolution & development 6(4): 219–226.

Mwinyi A, Bailly X, Bourlat SJ, et al. (2010) The phylogenetic position of Acoela as revealed by the complete mitochondrial genome of Symsagittifera roscoffensis. BMC evolutionary biology 10: 309.

Nielsen C (1995) Animal Evolution: Interrelationships of the Living Phyla. Oxford University Press.

Patro R, Duggal G, Love MI, et al. (2017) Salmon provides fast and bias-aware quantification of transcript expression. Nature methods 14(4): 417–419.

Pearson BJ, Eisenhoffer GT and Gurley KA (2009) Formaldehyde-based whole-mount in situ hybridization method for planarians. Developmental. Wiley Online Library. Available at: https://onlinelibrary.wiley.com/doi/abs/10.1002/dvdy.21849.

Perea-Atienza E, Sprecher SG and Martínez P (2018) Characterization of the bHLH family of transcriptional regulators in the acoel S. roscoffensis and their putative role in neurogenesis. EvoDevo 9: 8.

Peterson KJ and Eernisse DJ (2001) Animal phylogeny and the ancestry of bilaterians: inferences from morphology and 18S rDNA gene sequences. Evolution & development 3(3): 170–205.

Philippe H, Brinkmann H, Martinez P, et al. (2007) Acoel flatworms are not platyhelminthes: evidence from phylogenomics. PloS one 2(8): e717.

Philippe H, Brinkmann H, Copley RR, et al. (2011) Acoelomorph flatworms are deuterostomes related to Xenoturbella. Nature 470(7333): 255–258.

Philippe H, Poustka AJ, Chiodin M, et al. (2019) Mitigating Anticipated Effects of Systematic Errors Supports Sister-Group Relationship between Xenacoelomorpha and Ambulacraria. Current biology: CB 29(11): 1818–1826.e6.

Pimentel H, Bray NL, Puente S, et al. (2017) Differential analysis of RNA-seq incorporating quantification uncertainty. Nature methods 14(7): 687–690.

Ramachandra NB, Gates RD, Ladurner P, et al. (2002) Embryonic development in the primitive bilaterian Neochildia fusca: normal morphogenesis and isolation of POU genes Brn-1 and Brn-3. Development genes and evolution 212(2): 55–69.

Ruiz-Trillo I and Paps J (2016) Acoelomorpha: earliest branching bilaterians or deuterostomes? Organisms, diversity & evolution 16(2): 391–399.

Ruiz-Trillo I, Riutort M, Littlewood DT, et al. (1999) Acoel flatworms: earliest extant bilaterian Metazoans, not members of Platyhelminthes. Science 283(5409): 1919–1923.

Ruiz-Trillo I, Paps J, Loukota M, et al. (2002) A phylogenetic analysis of myosin heavy chain type II sequences corroborates that Acoela and Nemertodermatida are basal bilaterians. Proceedings of the National Academy of Sciences of the United States of America 99(17): 11246–11251.

Ruiz-Trillo I, Riutort M, Fourcade HM, et al. (2004) Mitochondrial genome data support the basal position of Acoelomorpha and the polyphyly of the Platyhelminthes. Molecular phylogenetics and evolution 33(2): 321–332.

Semmler H, Bailly X and Wanninger A (2008) Myogenesis in the basal bilaterian Symsagittifera roscoffensis (Acoela). Frontiers in zoology 5: 14.

Sempere LF, Martinez P, Cole C, et al. (2007) Phylogenetic distribution of microRNAs supports the basal position of acoel flatworms and the polyphyly of Platyhelminthes. Evolution & development 9(5): 409–415.

Sinigaglia C, Thiel D, Hejnol A, et al. (2018) A safer, urea-based in situ hybridization method improves detection of gene expression in diverse animal species. Developmental biology 434(1): 15–23.

Smith JPS, Teyler S and Rieger RM (1986) Is the Turbellaria polyphyletic? Hydrobiologia. DOI: 10.1007/bf00046223.

Srivastava M, Mazza-Curll KL, van Wolfswinkel JC, et al. (2014) Whole-body acoel regeneration is controlled by Wnt and Bmp-Admp signaling. Current biology: CB 24(10): 1107–1113.

Telford MJ, Lockyer AE, Cartwright-Finch C, et al. (2003) Combined large and small subunit ribosomal RNA phylogenies support a basal position of the acoelomorph flatworms. Proceedings. Biological sciences / The Royal Society 270(1519): 1077–1083.

